# Modified forms of easyCLIP

**DOI:** 10.1101/2021.12.15.472862

**Authors:** Douglas F Porter, Raghav M Garg, Robin M. Meyers, Weili Miao, Luca Ducoli, Brian J Zarnegar, Paul A Khavari

## Abstract

The easyCLIP protocol describes a method for both normal CLIP library construction and the absolute quantification of RNA cross-linking rates, data which could be usefully combined to analyze RNA-protein interactions. Using these cross-linking metrics, significant interactions could be defined relative to a set of random non-RBPs. The original easyCLIP protocol did not use index reads, required custom sequencing primers, and did not have an easily reproducible analysis workflow. This short paper attempts to amend these deficiencies. It also includes some additional technical experiments and investigates the usage of alternative adapters. The results here are intended to allow more options to easily perform and analyze easyCLIP.

## Introduction

The easyCLIP method, describing both a CLIP-seq library construction method (**Figure 1a**) and a UV protein-RNA cross-link rate measurement method, was presented in paper^1^ claiming the following strengths: (1) it was an easy, quick and reliable method to produce normal CLIP-seq libraries; (2) it provided troubleshooting information based on the fluorescent markers on adapters; (3) it quantified the absolute rate of protein-RNA cross-linking; (4) it enabled distinguishing RBPs from non-RBPs on the basis of combining cross-link rate and CLIP libraries; (5) on the same basis, it enabled distinguishing “specific” RNA interactions, *vs* those of a non-RBP.

**Figure 1.**
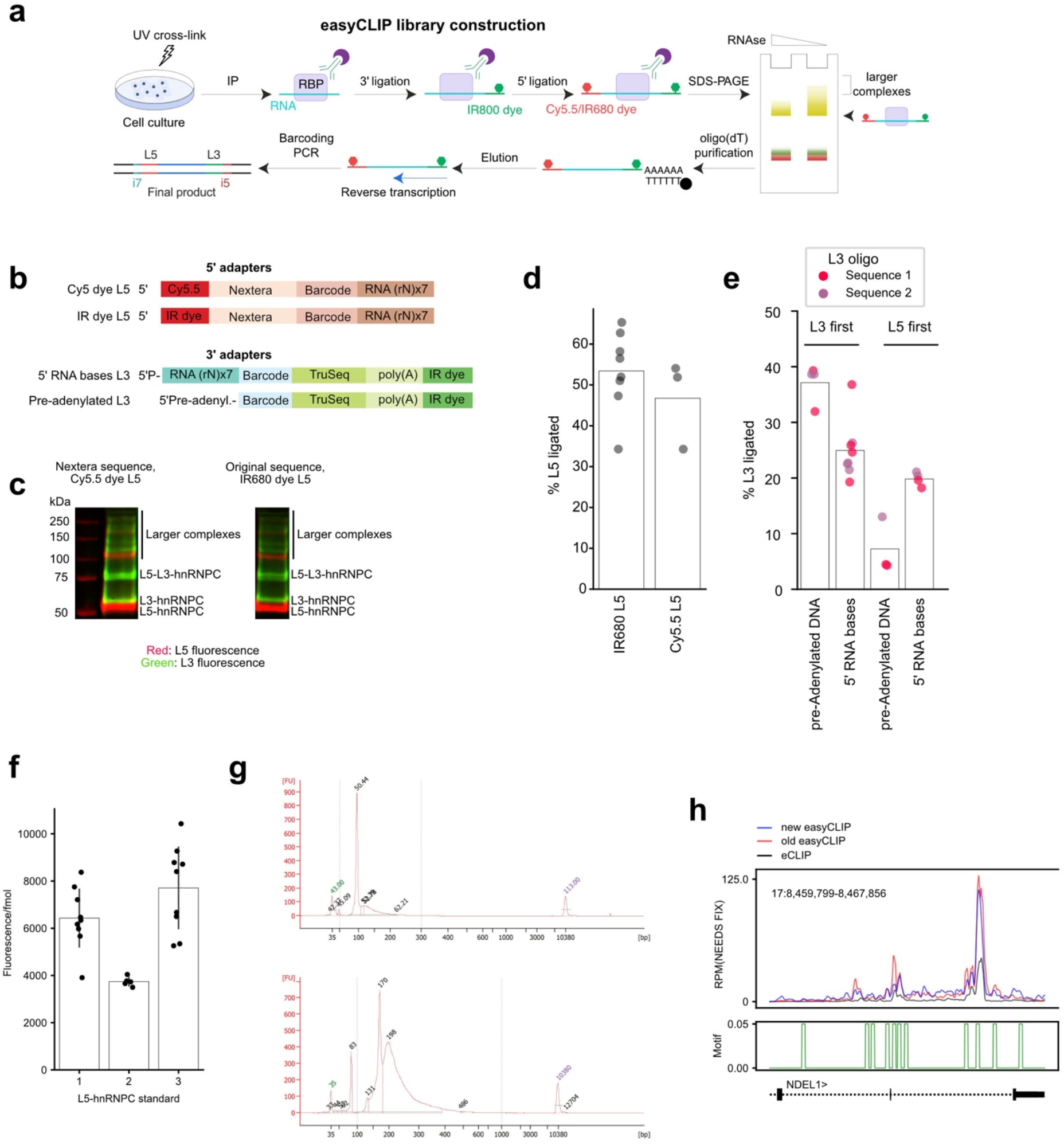
**a** The easyCLIP library preparation protocol. **b** Adapters for the easyCLIP protocol. The custom sequences in the original adapters are replaced here with Nextera (L5) and TruSeq (L3) sequences to enable standard PCR barcoding and sequencing primers. Two alternative dyes for L5 are shown, and two L3 5’ ends −5’ RNA bases or pre-adenylylated DNA (referred to as pre-adenylated in this paper). **c** An L5 oligo with the Nextera sequence and Cy5.5 dye generates a similar fluorescence shift from ligation, with similar brightness, as the original dye and sequence. Images are adapter fluorescence on nitrocellulose membranes of easyCLIP performed on hnRNP C. **d** The ligation efficiency of L5 oligos with Cy5.5 dyes or the original IR680 dyes, as assayed on hnRNP C, were similar. **e** The efficiency of L3 ligations on hnRNP C, in which the L3 ligation was performed either before (“L3 first”) or after (“L5 first”) the L5 ligation. When the L5 ligation was performed first, a single batch of L5 ligations was split into separate replicates before the L3 ligations. When the L3 ligation was performed first, both L3 and L5 ligations were performed separately for each replicate. **f** Fluorescence values of L5 per fmol across three separate L5 preparations and hnRNP C RNA standard preparations, measured in 2.5% aliquots across numerous experiments. That is, each dot represents a measure of fluorescence for a 2.5% aliquot of a standard, for three different standard preparations. **g** DNA bioanalyzer traces of CLIP PCR product before (*top*) and after (*bottom*) agarose gel size selection and an additional 5 rounds of PCR to add the full-length, barcoded Illumina adapters. The initial PCR had 15 cycles. **h** Trace of eCLIP, the original easyCLIP, and the updated easyCLIP signal for RBFOX2 at the NDEL1 locus. Instances of the shorter form of the RBFOX2 binding motif (GCAUG) are plotted as smoothed green lines below.

To briefly review that paper’s conclusions: the method suggested that controlling CLIP data to random non-RBPs might be advantageous for defining specific interactions, as opposed to the traditional methods of RNA inputs, no-epitope purifications, or no-control CLIP. The ability to measure absolute cross-link rates was employed to test the effect on cross-link rates of the most common 32 missense mutations across all human cancers in RBPs (29 proteins), finding increased cross-link rates for missense mutations in A1CF, KHDRBS2, and PCBP1. The A1CF and KHDRBS2 effects could be validated *in vitro*, while the L100P/Q missense mutations of PCBP1 rendered the protein insoluble *in vitro*. A mild decrease in DDX3X crosslinking with its most common missense mutation was also observed. Finally, expression of the L100P/Q PCBP1 mutant led to transcriptomic changes that could help explain an oncogenic effect of the mutation.

After publishing easyCLIP, it became clear that the use of a non-standard sequencing primer and the absence of PCR indexes were major drawbacks to the method. In addition, the large easyCLIP code repository was not presented in an easily reproducible package to perform the most basic tasks. We have attempted to correct these deficiencies in this work.

## Results

### Adapter sequence modifications

The previous easyCLIP L5 adapter required a custom sequencing primer and the L3 adapter was not compatible with Truseq barcoding. Both primers were redesigned, switching the L5 adapter sequence to a standard Nextera sequence and the L3 adapter to a standard Truseq sequence (**Figure 1b**). These adapters do not require custom sequencing primers and are compatible with normal PCR barcoding. The combination of Nextera and Truseq prevents the formation of adapter-adapter sequence complementarity that would occur if both ends were Truseq or Nextera. It also has the advantage of decreasing contamination from other high-throughput sequencing libraries being prepared in the same lab, since it will only PCR amplify with the mixed primer set. The Nextera L5 adapter ligated with the same ~50% efficiency as the previous easyCLIP adapter (**Figure 1c**). Standard PCR barcoding is useful because it enables a single set of L3 and L5 adapters to be re-used for multiple samples and sequenced in the same lane.

The original easyCLIP L5 adapter was purchased with an azide 5’ group that was reacted with DBCO-conjugated IR680 dye. The IR680 modification is not (as of Septemer 2021) available for RNA oligos from IDT, but the Cy5.5 modification is available, so oligos using the Cy5.5 dye substituted for IR680 were purchased. The Cy5.5 dye could be used to visualize the L5 adapter on nitrocellulose with similar brightness as the IR680 dye in the 700 nm channel on an Odyssey scanner (**Figure 1c**) and did not impair ligation efficiency (**Figure 1d**). This modification permits the usage of purchased L5 adapters without any modifications required.

### L3 Adapter ligation efficiency

If the L5 adapter ligation performed at ~50%, up to ~100%, while the L3 ligation, which uses pre-adenylylated 5’ ends (often termed 5’adenylated or pre-adenylated), tended to perform more poorly, it seemed possible the L3 ligation efficiency could potentially be boosted by mimicking the L5 ligation conditions – specifically, by adding 5’ RNA bases to the L3 adapter.

To review the relevant mechanism, T4 RNA ligase joins 5’PO_4_ and 3’OH from “donor” and “acceptor” (respectively) polynucleotides by (1) initially reacting with ATP to form a covalent bond with AMP, (2) transferring covalently bound AMP to the 5’PO_4_ group of the donor strand to create an adenosine group connected through a double phosphate bond to the rest of the donor strand, and (3) displacement of AMP by the 3’OH of the acceptor strand to form a completed ligation^2^. In a crystal structure, T4 RNA ligase makes RNA contacts with the 3’OH-providing substrate, but does not contact the RNA bases of the 5’PO_4_-providing substrate^3^, consistent with its requirement for RNA bases in the 3’OH-providing substrate and not in the 5’PO_4_-providing substrate^4^. RNA ligase can also pre-adenylylate DNA oligos efficiently as long as the 5’ base is not G^5^. These facts suggest the pre-adenylylated DNA oligo would be competitive with the 5’RNA bases form, but do not prove either would necessarily work better in the context of CLIP.

Testing ligation efficiencies of pre-adenylylated or 5’RNA base oligos in CLIP-seq led to mixed results, with variations in ligation efficiency between experiments and depending on whether an L5 ligation was performed first (**Figure 1e**). The modest increase in efficiency for pre-adenylylated L3 when the L3 ligation is performed first may reflect the avoidance of other possible competing ligation products (including circularization) that may form with ATP present in the ligation. RNAse I, which is used in this protocol, leaves 5’OH ends, which should not permit circularization (T4 RNA ligase lacks polynucleotide kinase activity), although polynucleotide kinase remaining on the beads from the 3’ dephosphorylation step could produce 5’ phosphorylated ends — suggesting more stringent washing might improve efficiency. After the L5 ligation, circularization is not possible, and other competing ligation products may be slower to form in a second ligation. As a result, the pre-adenylated form might lose this advantage when the L5 ligation is performed first. Why the non-pre-adenylylated form was more efficient in this case is not clear, but RNA substrates may be more efficient in this context.

### Adapter synthesis

The RNA-DNA hybrid L3 also requires no modification of the oligo besides labelling with dye. A summary of costs of oligo synthesis (from IDT) and modification is given in **Table 1**. The dye modifications are of course not necessary if only library preparation is required, and if no troubleshooting information is needed. If separate PCR barcodes were always used, no in-line barcodes would be necessary, and a single L5 and L3 could be used for all samples. However, it is helpful to have a few different L5/L3s to enable sample pooling during the CLIP process.

**Table 1.**
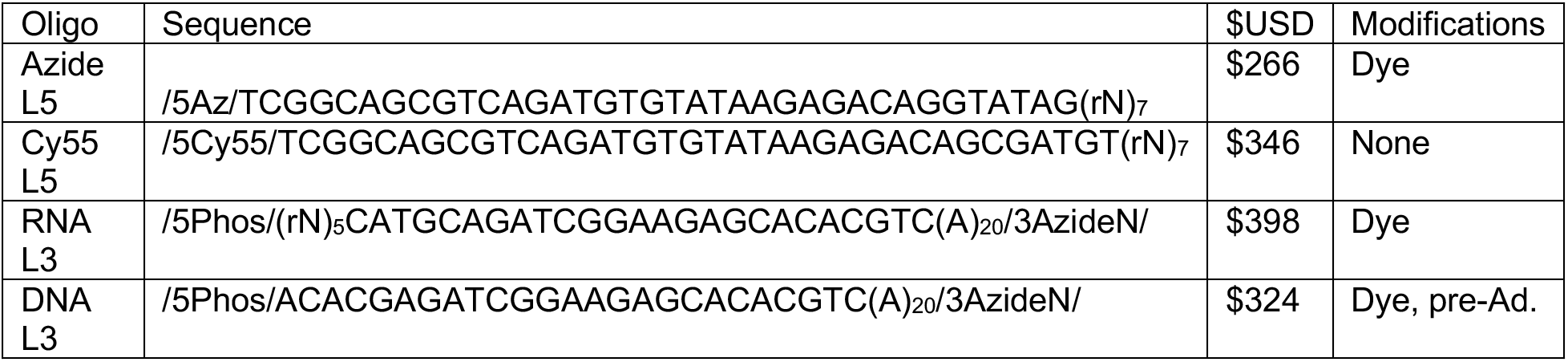
Costs of oligos for CLIP-seq. Costs are listed were obtained from the IDT website on 9/25/2021, for 100 nmol oligos with an RNAse-free HPLC purification. Costs are in US dollars. The modifications column denotes modifications not performed by IDT. 100 nmol scale is sufficient for IDT to generally produce several nmol of oligo, which is enough adapter for hundreds of CLIP-seq experiments. /5Az/, 5’ azido group. /5Cy5/, 5’ Cy5.5 dye. /5Phos/, 5’ phosphorylation. /3AzideN/, 3’ azido group. rN, random RNA base.

### RNA fluorescence standards

Creating and running aliquots of a standard preparation of quantified RNA standard adds some difficulty to the easyCLIP procedure. To determine if it was always necessary to include such a standard, cross-link rate determination experiments in easyCLIP were reanalyzed for consistency in the standard aliquots. Fluorescence per oligo differed for different L5 oligo preparations, indicating that quantifying a standard at least once is necessary (**Figure 1f**). Across 23 gels, the fluorescence between experiments was always within 2-fold of the mean for the same standard, and within 50% of the mean in 22/23 cases (**Figure 1f**). Variability in aliquoting, boiling and gel loading could all increase variation, and might account for much observed disparity. These results indicate that if a standard is not run, but has a known mean signal, estimates are likely to be accurate to within ~50%.

### Sequencing

A common strategy in sequencing library construction is to perform an initial PCR, size select libraries using a gel, and then perform an additional PCR with barcoding primers. Alternatively, a SPRI bead clean-up may be done between primary and secondary PCRs, with the gel performed at the end. In CLIP, a gel purification is generally performed to remove no-insert adapter-ligations and PCR product.

CLIP-seq libraries using the modified oligos and library preparation protocol were prepared for three replicates of RBFOX2 in 293T cells. An initial PCR with un-indexed primers was performed on the reverse transcription product. To perform a clean-up and concentration of the product, polynomial curves were fit to the yield-vs-isopropanol-vs-length numbers in Table 1 of Fishman *et al*.^6^, and isopropanol concentrations were linearly extrapolated past the tested conditions to determine a reasonable isopropanol and SPRIselect bead to preserve CLIP PCR product while reducing primers. After SPRIselect clean-up, samples were then run on a 4% agarose gel. Signal above the linker-dimer band was extracted and input to a second PCR reaction with primers indexed according to the Illumina dual-indexing scheme. Despite the low resolution of agarose gels, this quick gel extraction was successful in reducing linker dimers (**Figure 1g**). The dual ligated RNA could be amplified and sequenced as with the original adapters and the sequencing trace for RBFOX2 at *NDEL1* was quite similar to previous easyCLIP and eCLIP data (**Figure 1h**). To ensure a PCR product is visible in a gel, the gel is probably more reliably performed after both PCRs.

### Improvements to easyCLIP processing pipeline

The easyCLIP processing pipeline was modified to improve its accessibility and simplicity. The pipeline is now implemented with a standardized snakemake workflow, conda environment and configuration system (https://github.com/dfporter/easyclip-snakemake). Processed data files are now available on-line (http://khavarilab.stanford.edu/datasets-1) and the random non-RBP control protein datasets are downloaded automatically in the pipeline. Results with this new pipeline were highly similar to our previous report (**Figure 2**).

**Figure 2.**
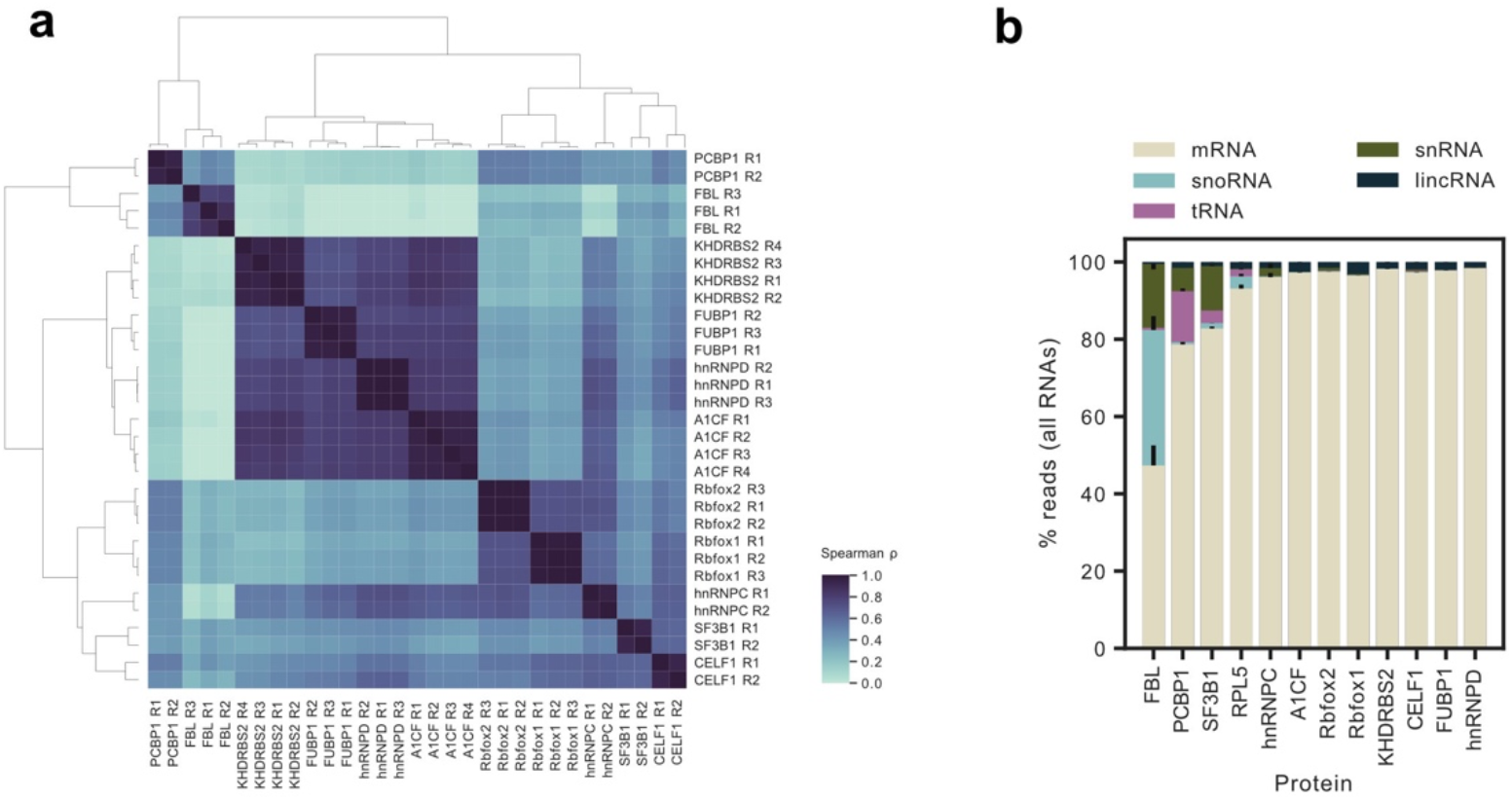
Similar results were obtained with the new snakemake workflow as the previous workflow. These figures re-analyze the data in Porter *et al*. and are comparable to Figure 2D and 2G in that work. **a** Spearman ρ values for reads-per-gene from replicates of 11 RBPs, excluding RPL5 as it has only one definitive target RNA (5S rRNA), using the datasets obtained previously. Endogenous hnRNP C, RBFOX2 and FBL were immunopurified, HA-PCBP1 was stably integrated, and the others were transiently expressed as HA-tagged forms. PCBP1 was expressed in HCT116 cells and the others in 293Ts. **b** Fraction of easyCLIP reads for of the indicated RBP mapping to RNAs of the given ENSEMBL biotype. Bars represent the standard deviation.

## Discussion

This paper presents some possible modifications to the easyCLIP protocol. Pre-adenylation *vs* the inclusion of RNA bases in the L3 adapter were compared, which suggested both are feasible options, with the 5’RNA bases oligos having more consistent ligation efficiencies, and the pre-adenylated DNA oligos performing better when used as the first ligation. The software pipeline was simplified and streamlined, with additional documentation. The results here are intended to allow more options to easily perform and analyze easyCLIP.

## Methods

### CLIP

The CLIP method is described in Table S1. A few adjustments may be noted: (1) the protocol includes the option of the 5’phosphorylated linker; (2) linker and primer sequences have been changed; (3) a low salt wash is no longer included; (4) a new tab describing PCR indexing and gel clean-up is included; (5) a general-purpose RNAse concentration suggested. Note that if not using pre-adenylylated linkers, the RNAse used should be one that, like RNAse I, does not leave 5’ phosphate groups.

### Linker labelling

Linkers were resuspended in 70 μL PBS or water. 700 μL Qiagen buffer PNI and 210 μL isopropanol (100%) were added to the mixture, vortexed, and loaded onto two nucleotide removal columns (Qiagen). Qiagen buffer PNI can be replaced with a mixture of 1 volume Qiagen buffer PB to 0.75 volumes isopropanol. Columns were centrifuged at 6,000 RPM for 1 minute, flow-through discarded, the remaining oligo mixture added to the columns, and centrifuged again. The flow-through was discarded, 750 μL 85% ethanol was added to the column, and columns centrifuged 6,000 RPM for 1 minute. This wash was repeated. After washing, the columns were centrifuged for 1 minute at 17.9 krcf to dry the columns, then columns were moved to Eppendorf tubes and 50 μL water added to each column; the tip of the pipette can touch the center of the column to ensure liquid is delivered the center of the column. The columns were left with water for 1 minute before centrifuging to elute. The concentration was determined using A260 values. An equal volume of RNAse-free PBS buffer was added to the eluates.

IRDye 680RD DBCO (0.5 mg) (LI-COR, 429 nmol) was resuspended in 42.9 μL phosphate-buffered saline (PBS) yielding a 10 mM dye mixture. 4-5 μL of 10 mM dye (40-50 nmol) was added to 10–150 μg purified oligonucleotide (~1–12 nmol) in PBS for a total volume of 200 μL and incubated for 2 h at 37°. Labelled oligos were purified again using the same Qiagen nucleotide removal kit procedure, eluting in water, and adding at least an equal volume of PBS before diluting, aliquoting and freezing.

## Supplementary tables

**Table S1.** The modified easyCLIP protocol, reagents and oligonucleotides.

## Data availability

The example RBFOX2 data has GEO accession GSE190678.

## Acknowledgments

This work was supported by a USVA Merit Review grant BX001409 to P.A.K. and by NIAMS/NIH grants AR49737 and AR45192 to P.A.K., 1F32AR072504 to D.F.P., and K01AR071481 to B.J.Z. We thank Babette Hammerling of Illumina for the suggestion of mixing the Nextera and Truseq sequences.

